# Improved Activity and Kinetics of Endoglucanase Biofuel Enzyme with Addition of an AzoTAB Surfactant

**DOI:** 10.1101/2023.08.20.554048

**Authors:** Zumra Peksaglam Seidel, C. Ted Lee

**Affiliations:** Department of Chemical Engineering and Materials Science, University of Southern California, Los Angeles, California 90089 U.S.A

**Author notes:** Corresponding author: Department of Chemical Engineering and Materials Science, University of Southern California, Los Angeles, CA 90089-1211. E-mail addresses and.

**Keywords:** photosurfactant, azobenzene-based surfactants, enzyme surfactant interaction, cellulase enzymes, surfactant-added heterogenous reaction

## Abstract

Endoglucanases degrade *β*-1,4-glycosidic bonds of crystalline cellulose into insoluble or soluble cellooligosaccharides at the solid-liquid interface. The enhanced activity and kinetics of endoglucanase from *Aspergillus niger* with addition of a light responsive azobenzene trimethyl ammonium bromide (azoTAB) surfactant is studied. AzoTAB is a photoresponsive surfactant that exists as a relatively-hydrophobic *trans* isomer under visible light (434 nm) and a relatively-hydrophilic *cis* isomer under UV light (350 nm). Endoglucanase catalytic activity can be controlled with light illumination slightly for microcrystalline cellulose natural substate (∼15%) and significantly for p-nitrophenol-based model substrate (∼2-fold) by switching between the *trans* (higher enzyme binding affinity, resulting higher enzyme unfolding) and *cis* form of azoTAB. Endoglucanase activity increases 45% towards avicel crystalline substate and 4-fold towards 4-nitrophenyl β-D-cellobioside substrate in the presence of 0.4 mM azoTAB under UV light (90% *cis* and 10% *trans* isomers). In comparison, endoglucanase catalytic activity increases 5-10% towards crystalline cellulose substrate with the addition of sodium dodecyl benzene sulfonate (SDBS), sodium dodecyl sulfate (SDS) and dodecyl trimethyl ammonium (DTAB). 0.4 mM AzoTAB-UV addition leads to an increase of maximum endoglucanase adsorption (*E*_max_) from 7.89 mg enzyme/g avicel to 12.92 mg enzyme/g avicel and catalytic enzyme efficiency (*k*_cat_/*K*_M_) from 0.031 L/(mg.s) to 0.061 L/(mg.s). It is found that adsorbed enzyme concentration on substrate correlates to enzyme specific activity for azoTAB containing reaction. Additionally, 40-50% activity enhancement and increased bound enzyme to substrate are detected with azoTAB addition at different enzyme, substrate, and inhibitor concentrations. Improvement of substrate properties of azoTAB addition is associated with lower reciprocal terms of Michaelis constant (*K*_M_), the adsorption coefficient (*K*_ad_) and fractal parameter (*h*) values similar to the presence of other surfactants in this study and the literature. Furthermore, 45-50% activity enhancement via azoTAB surfactant preserved for all three cellulase enzyme mixture of endoglucanase, cellohydrolase and *β*-glucosidase. Consequently, azoTAB can be applied as profitable additive for the heterogenous enzymatic cellulose hydrolysis, resulting in a 30% decrease in the enzyme load based on the specific activity, adsorption, and fractal kinetics results.

## Introduction

Cellulose, a homopolymer of *β*-1,4-linked glucose monomers, is the main constituent of plant cell walls and one of the primary polysaccharides in biomass.^1^ Cellulose can be hydrolyzed by the cellulase multienzyme complex into fermentable sugars (i.e., glucose) and hence fermented into bioethanol as a readily-available sustainable energy source. Enzymatic saccharification of cellulose into glucose requires three enzymes: random breakage of internal glycoside bonds by endoglucanase (endocellulase, *β*-1,4-endoglucan hydrolase, *β*-1,4-*β*-D-glucanase, EC 3.2.1.4), cellobiose (i.e., two linked glucose molecules) cleavage from the terminal ends of cellulose chains via cellobiohydrolase (exocellulase, EC 3.2.1.91), and hydrolysis of cellobiose and short-chain cellooligosaccharides into fermentable glucose by *β*-glucosidase (cellobiase, EC 3.2.1.21).

One of the major challenges of the above process is the high cost of saccharification enzymes due to the need for high enzyme concentrations combined with irreversible enzyme-substrate binding and slow enzymatic cellulose hydrolysis process.^2^ The catalytic activity of cellulase enzymes decreases quickly as a result of this nonproductive binding of enzymes to crystalline substrate and end-product inhibition.^3^ The enzymes bound to the substrate surface cannot be recovered and hence high concentrations of enzymes are required. As an illustration, 30 grams of enzyme is needed per liter of bioethanol.^4^

The enzymatic conversion of cellulose is a heterogeneous reaction that occurs at solid-liquid interface between soluble enzymes and crystalline cellulose. Higher fermentable sugars can be obtained at lower enzyme loading with surfactant addition.^5^ Easy access of enzymes to the active sites of cellulose and prevention of inactivated adsorbed enzymes have been enhanced the cellulose hydrolysis process yield by upon surfactant addition.^6, 7^ The adsorption of surfactants on substrate surfaces lowers irreversible enzyme binding to the substrate. Additionally, enzyme-surfactant interaction minimizes enzyme denaturation and increases enzyme stability.^8^

In this work, improvement of endoglucanase activity and kinetics is studied with the addition of a photoresponsive surfactant, azobenzene trimethylammonium bromide (azoTAB) that undergoes photoisomerization through nitrogen double bond rotation upon exposure to visible and UV light. AzoTAB primarily exists as the *trans* isomer (75/25 *trans*/*cis*) with planar structure and lower dipole moment of the nitrogen bond under visible light (434 nm), whereas under UV light (350 nm) the *cis* form (10/90 *trans*/*cis*) of azoTAB is predominant (see Scheme 1).^9^ The lower dipole moment of the *trans* photoisomer leads to the *trans* form of azoTAB being relatively more hydrophobic compared to the *cis* isomer. The more hydrophobic *trans* isomer of azoTAB has a greater tendency to bind and unfold proteins compared to the *cis* form due to the hydrophobicity differences of the photoisomers. The reversible hydrophobicity of the surfactant upon light exposure has been applied to control protein unfolding, activity, and dynamics.^10–13^ In our earlier study, *β*-glucosidase and azoTAB interaction leads to *dimer-to-monomer* transition associated with 50% activity enhancement of *β*-glucosidase towards cellobiose natural substrate.^14^ In this study, we explore endoglucanase enzyme activity and kinetics in the presence of azoTAB. All in all, azoTAB can be used as an additive to lower enzyme saccharification cost of bioethanol production.

**Scheme 1.**
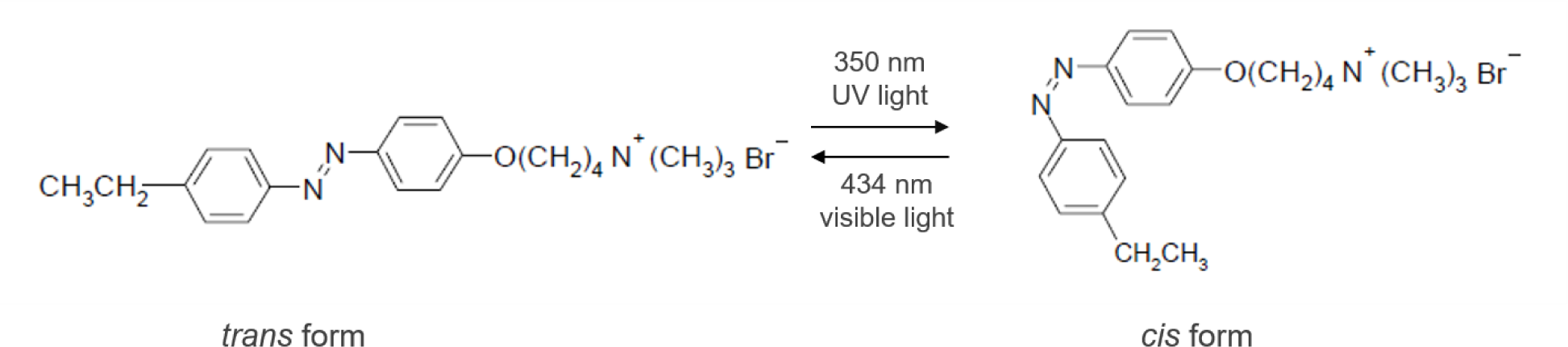
Molecular structure of 4-ethyl-4’(trimethylamine-butoxy) azobenzene bromide (azoTAB) photo-surfactant

## Experimental Section

### Materials

The cationic azobenzene trimethyl ammonium bromide surfactant (*M*_W_=420 g/mol) shown in Scheme 1 was synthesized according to previous studies.^9, 15^ Briefly, 4-ethylaniline was azo-coupled with phenol; subsequently alkylated by 1,4-dibromobutane and followed by quaternization with trimethylamine. All chemicals were purchased from Millipore Sigma at the highest purity unless otherwise stated. The *trans*-to-*cis* photoisomerization of azoTAB was obtained with an 84-W long-wave (365 nm) UV lamp (Spectroline, Model no. XX-15A). The experiments of UV light samples were performed in the dark due to the fact that the thermal conversion of *cis*-to-*trans* azoTAB is ∼24 hours in the dark at 25 °C.

### Purification of endoglucanase

Cellulase from *Aspergillus niger* was acquired from Sigma-Aldrich (Cat. no. 22178). 4 grams of the off-white crude powder was dissolved in 100 mL of protein buffer (25 mM sodium phosphate buffer pH 6.5, 50 mM NaCl, 10% glycerol) resulting in 40 mg/mL of protein solution. The protein solution was filtered and applied to a 5 mL HiTrap Q HP anion exchange column (GE Healthcare, Cat. no. 17115301) and later washed with 30× column volume with the protein buffer. The protein was eluted at a flow rate of 2 mL/min by salt gradient against the elution buffer (25 mM sodium phosphate buffer pH6.5, 550 mM NaCl, 10% glycerol) using fast protein liquid chromatography, FPLC (ÄKTA Pure from GE Healthcare) at 4 °C and the eluted fractions were run on SDS-PAGE. All fractions were tested using Nanodrop 2000 (Thermo Scientific, USA) at 280 nm to detect the presence of protein and provide a rough estimate of protein concentration. Equal concentrations of fractions were applied for SDS-PAGE and measured enzyme activity against avicel microcrystalline substate to identify endoglucanase fraction. The active fractions were pooled, concentrated using Amicon Ultra 4 mL centrifuge tubes with 10 kDa cutoff membranes (UFC801024), and concentrated protein solutions were kept at 4 °C for further studies.

### Measurement of Endoglucanase Activity

Endoglucanase activity was detected with Somogyi-Nelson procedure,^16^ by measuring the reduced ends of avicel, a natural microcrystalline cellulose, upon conversion to insoluble and soluble cellooligosaccharides. Briefly, the reaction took place in 2 mL vials (1.1 mL reaction volume) at a final concentrations of 0.01 mg/mL (0.001%, w/v) avicel and 100 *μ*g/mL purified endoglucanase fraction and various azoTAB concentrations in 50 mM sodium acetate buffer, pH 5 at 25 °C. The reaction vial was constantly stirred at 200 rpm with a magnetic stir bar to prevent microcrystalline cellulose settling at the bottom of the vial. When desired, azoTAB was pre-converted to the *cis* state under UV light and then mixed into the reaction solution with the reaction performed at the dark to minimize photo-transition to *trans* isomer. At the initial time (*t*_0_) and every ten minutes for 90 minutes, 50 *μ*L of reaction mixture was diluted ten-fold and stopped with 50 mM phosphate buffer, pH 9.5. This stopped sample solution was combined with 500 *μ*L of alkaline copper tartrate and heated for 20 minutes in boiling water to induce cuprous oxide with the reduced ends of avicel. Thereafter, 500 *μ*L of arsenomolybdic acid was added into the samples to allow molybdenum blue to form for five minutes at room temperature from the interaction between molybdic acid and cuprous oxide. The resultant solutions were diluted eleven-fold in deionized water and the absorbance at 760 nm was monitored with UV-vis spectroscopy (Agilent, model 8453). Similar to our previous study,^14^ the absorbance at 760 nm was background corrected by subtracting the absorbance at 500 nm, where the absorbance of both azoTAB and molybdenum blue is essentially zero. The endoglucanase catalytic activity was determined from the initial rate increase in the corrected absorbance applying data fitting to (*A*_t_-*A*_∞_)/(*A*_0_-*A*_∞_) =e^-at^, where *A*_t_ is the absorbance at time *t*. The optimum surfactant concentrations (0.4 mM for azoTAB under visible and UV light; 1.5 mM for SDS, SDBS and DTAB) were used for the further studies unless otherwise stated.

The enzyme catalytic rate kinetic constants, *k*_cat_ and *K*_M_ were calculated using 0.001-40 mg/mL avicel with 100 *μ*g/mL endoglucanase. The nonlinear regression model with Michaelis-Menten equation was applied, *V*_max_ and *K*_M_ values were calculated by minimizing the residual sum of squared errors method. For comparison, Hanes-Woolf linear equation was used to calculate *V*_max_ and *K*_M_.

### Photoreversible Endoglucanase Activity

Light responsive endoglucanase activity was performed towards 4-nitrophenyl *β*-D-cellobioside model substrate (Millipore Sigma-N5759) under visible and UV light. The reaction took place at a final concentration of 5 mM substrate, 0.3 mg/mL enzyme and 0.4 mM pre-converted azoTAB solution in 50 mM sodium acetate (pH 5) buffer at 37 °C. Every 2 minutes, 500 *µ*L of the reaction sample was stopped into 500 *µ*L of 50 mM phosphate (pH 9.5) buffer. The p-nitrophenol product (p-NP) was monitored at 429 nm with extinction coefficient of 13.2 mM^-1^cm^-1^ to avoid strong absorption peak effect of azoTAB similar to our previous study.^14^ Subsequent to 10 minutes of reaction under visible or UV light, the light conditions were switched to record light-induced enzyme catalytic rate.

### Optical microscopy

Final concentrations of 0.5 mg/mL endoglucanase, 0.03 mg/mL avicel were mixed in 8 mL screwcap tubes and filled with 5 mL solution in 50 mM sodium acetate, pH 5 buffer with and without 0.4 mM azoTAB and 200 mM glucose. The reaction vials were mixed at 240 rpm for six days. 300 *µ*L reaction samples were taken at time 0 min, 3^rd^ and 6^th^ hour, 2^nd^, 4^th^ and 6^th^ days and terminated into 300 *µ*L 50 mM phosphate (pH 9.5) buffer. The samples were observed using an Olympus IX71 inverted microscope equipped with a 10× objective lens. Optical microscopy images were captured with a CCD digital camera (Hamamatsu, model no. C4742-95).

### Endoglucanase Adsorption on Avicel

Twenty-two milligrams of avicel (2 wt%) were placed in 2-mL screwcap tubes and filled with 1.1 mL of reaction solution in 50 mM sodium acetate buffer at pH 5 with a total endoglucanase concentration between 0.1-2 mg/mL for adsorption assays. The optimum activity associated surfactant concentrations were selected: 0.4 mM azoTAB visible and UV; 1.5 mM SDS, SDBS and DTAB. The reaction vials were stirred at 200 rpm with a magnetic stir bar for 40 minutes to allow the enzyme to bind to the substrate surface. Thereafter, 400 *µ*L reaction sample was taken and centrifuged at 13,000 rpm for a couple of seconds to settle down microcrystalline substrate. The enzyme content in the supernatant was measured at 280 nm with a subtraction of 600 nm wavelength using UV-vis spectroscopy. The same procedure including magnetic stir bars without the substrate were repeated to calculate calibration lines for pure and surfactant added solutions separately for each condition due to 350nm strong wavelength of azoTAB and 260 nm peak of SDBS. The adsorbed endoglucanase was determined from the difference between the initial enzyme concentration and the calculated supernatant concentration value. Simply, the calibration lines were drawn for each condition: pure, azoTAB-vis, azoTAB-UV, SDS, SDBS and DTAB. The measured data points were fitted into each calibration lines with simple ordinary linear equation (i.e., y=mx+b) to find hydrolysate liquid phase enzyme concentration. These values corresponds to free enzyme concentration. The difference between the initial enzyme concentration and the free enzyme concentration was calculated to find adsorbed enzyme concentrations. Endoglucanase adsorption experiments were conducted with four replicates to acquire the adsorption isotherms.

Langmuir isotherm has been widely used cellulase adsorption model to determine adsorption parameters with the assumptions of a monolayer and reversible adsorption on cellulose binding sites without interactions with other enzymes. The enzyme adsorption was assumed to follow Langmuir isotherm as in equation (1)^17, 18^:

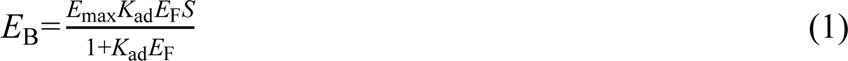

where *E*_B_ is the adsorbed enzyme concentration (mg/mL), *E*_max_ is the maximum adsorbed enzyme concentration (mg enzyme/g substrate), *E*_F_ is the free enzyme in solution (mg/mL), *S* is the substrate concentration (g/mL) and *K*_ad_ is the adsorption coefficient. *E*_max_ and *K*_ad_ were interpreted with linearized form of equation (1)^17, 19^:

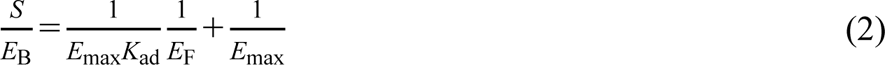

with *S*/*E*_B_ on the y-axis and 1/*E*_F_ on the x-axis.

The same experiment protocol with a constant endoglucanase concentration of 1 mg/mL and substrate concentration of 20 mg/mL was used to measure end-product inhibition effect with 0-200 mM of glucose or cellobiose with and without surfactant solutions. The same solutions without substrates were run in parallel and the average absorbance of them was used as initial absorbances.

The difference between the supernatant of glucose or cellobiose added samples and initial absorbance of control solutions was determined to find surfactant effect on the adsorbed enzyme concentrations as a function of glucose or cellobiose end-product concentration.

### Fractal kinetics

A final concentration of 0.01 mg/mL avicel and 62.5 *µ*g/mL endoglucanase were mixed with and without surfactant solutions to calculate fractal exponent, *h* and rate constant, *k* were determined using equation (3)^20, 21^:

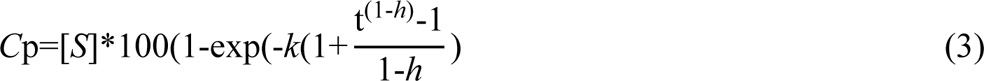

where *C*_p_ was the product concentration and [*S*] was the substate concentration (10 mg/L). Experimental product concentrations were calculated with the measured glucose extinction coefficient of 0.24744 L mg^-1^cm^-1^ at 760 nm minus 500 nm wavelength. Residual sum of squared errors method was applied to quantify *h* and *k*.

### Surfactant effect on enzyme mixtures

Using combination of enzymes, the hydrolysis rate was monitored in the absence and presence of surfactants.

Glucosidase from Aspergillus niger (Sigma-Aldrich cat. no. 49291) was purchased as a source of *β*-glucosidase and was purified as previously reported.^14^ Briefly, the brown powder solubilized at pH 7.5 and saturated with 80% ammonium sulfate overnight at 4°C. Then, the protein precipitate was washed 3 times with ammonium sulfate solution and resuspended in 50 mM sodium phosphate (pH 7.2) buffer including 1 M ammonium sulfate. Subsequently, HiTrap Phenyl HP (GE Healthcare, 17-1351-01) hydrophobic column was used to obtain *β*-glucosidase fraction. The fraction was desalted using 50 kDa membrane centrifuge tubes.

Cellobiohydrolase from Hyrpocrea jerorina (Trichoderma reesei) (Sigma-Aldrich cat. no. E6412) which is the most common component of enzyme mixtures for cellulose hydrolysis,^22,23^ was aliquoted and kept at −20 °C until usage. Enzyme solution was washed in 10 kDa membrane centrifuge tubes 4-5 times with 50 mM sodium acetate pH 5 buffer before adding to reaction tubes.

Optimal *β*-glucosidase and cellobiohydrolase enzyme ratios were selected and final enzyme concentrations of 60 *μ*g/mL of cellobiohydrolase, 62.5 *μ*g/mL endoglucanase and 30 *μ*g/mL *β*-glucosidase were used towards 0.01 mg/mL avicel. The surfactant effects on initial rate and long-term behaviors were examined with every 10 minutes collected samples for the first hour and two samples at times 5^th^ hour and 24^th^ hour reactions, respectively. Somogyi-Nelson method was applied to measure the samples similar to endoglucanase activity measurement. The rate was calculated with data fitting to four replicates and the difference between initial time (*t*_0_) to 1^st^ hour, 5^th^ hour and 24^th^ hour.

## Results and Discussion

Endoglucanase from *Aspergillus niger*, the most common industrial fungi,^24^ is purified from commercially available cellulase enzyme. The crude enzyme contains at a minimum of seven protein fractions as shown in Figure S1 (lane 2). The dissolved crude enzyme solution is applied into strong anion exchange column to separate the endoglucanase enzyme fraction from the other fractions. Positively-charged proteins are expected to bind to the anion exchange column strongly, while negatively-charged proteins and impurities would pass through the column without binding. According to SDS-PAGE, the flow through does not contain any protein fraction and further shows no activity against avicel substrate. Yet surprisingly none of the purified protein fractions display endoglucanase activity without the addition of flow through, while returning and mixing the flow through with all of the purified protein fractions at a ratio between 20/80 an 80/20 show endoglucanase catalytic activity. All purified fractions are catalytically active after mixing with flow through (150 mL flow through in 1100 mL reaction mixture). The flow through itself and mixture of bovine serum albumin, lysozyme and *β*-glucosidase three different enzymes do not show any endoglucase catalytic activity.

Endoglucanases from *Aspergillus niger* species are reported to have various isozymes having identical catalytic functions with divergent molecular weights.^25, 26^ Commercial enzyme preparations combine these diverse isoenzymes to provide diverse associations with the insoluble substrate, thus, break glycosidic bonds more efficiently. We speculate that the flow through of the anion exchange column possibly contains previously unreported cofactors of endoglucanase that keep the enzymes active. Interestingly, prior endoglucanase purification studies do not mention a loss of the catalytic activity of the purified fractions in the absence of cofactors in the literature. The major protein fraction (with ImageJ calculation: 35%) with a molecular weight of 48kDa was chosen, pooled, concentrated and used for further studies.

As the surfactants are more cost-efficient than the biofuel enzymes,^27^ the most favorable surfactant concentration is studied. The catalytic activity of purified endoglucanase is measured using avicel substrate as a function of azoTAB concentration under visible and UV light in Figure 1. While the concentration of azoTAB initially increases until 0.4 mM, the endoglucanase activity enhances to a maximum of 30% and 45% activity enhancement under visible and UV light, respectively. Although azoTAB surfactant photoisomers have a different hydrophobicity and hence binding affinity towards proteins, photoresponsive endoglucanase relative activity changes are minute. Similarly, the increase of surfactant tail length from C12 to C14 or C16 shows minimal activity changes on cellulose hydrolysis with endoglucanase enzyme.^28^ Additional increases in azoTAB concentration, however, led to a decrease in endoglucanase catalytic activity. Similar to our previous work with *β*-glucosidase enzyme,^14^ ∼50% activity enhancement is observed for the purified endoglucanase with azoTAB surfactant addition under UV light.

**Figure 1.**
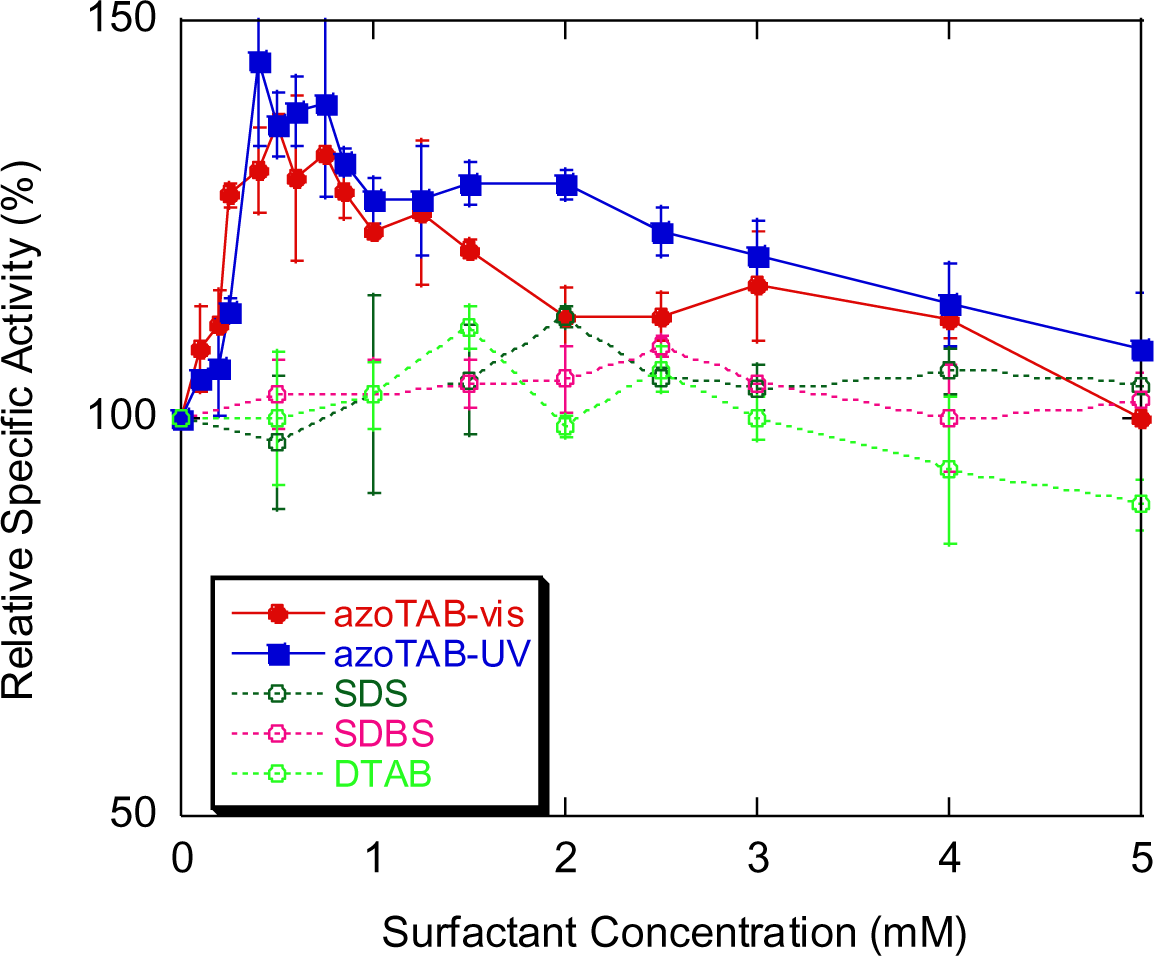
Relative specific activity of endoglucanase (100 *μ*g/mL) as a function of azoTAB (under visible or UV light), SDS, SDBS and DTAB concentration towards 0.01 mg/mL avicel. pH= 5, T= 25 °C. Lines are drawn to guide the eyes.

For comparison, SDS (sodium dodecyl sulfate), SDBS (sodium dodecyl benzene sulfonate; benzene containing frequently used surfactant) and DTAB (dodecyl trimethyl ammonium bromide; no-stimuli responsive, straight hydrophobic tail analog of azoTAB) surfactants are applied to compare benzene and TAB (trimethyl ammonium bromide) containing surfactants on endoglucanase activity. Maximum of 5-12% endoglucanase activity increase is observed at relatively higher concentrations (i.e. 1.5-2 mM common surfactant concentration versus 0.4 mM azoTAB concentration) of these traditional surfactants. Similar to our study, SDS addition increased the endoglucanase activity at low concentrations and above 2 mM addition of the surfactant addition reduces the catalytic activity of endoglucanase.^29^ The 5-12% increase of catalytic activity could be associated with the typical surfactant application on cellulose hydrolysis due to minimization irreversible substrate-enzyme binding.^6^ Similarly, azoTAB addition shows 20-30% of endoglucanase catalytic activity enhancement at the 1.5-2 mM surfactant concentration range. In literature, Tween 20 (35%),^30^ Tween 80 (27%-34%)^31, 32^and Triton X-100 (23%)^31^ chemical surfactants have been reported to result in 23-35% endoglucanase activity increases.

Additionally, surfactant and avicel substrate are incubated overnight before adding endoglucanase to understand preincubation effect on endoglucanase activity. Avicel contains a homogenous ∼50 µm particles (according to the manufacturer) of parallel glucose chains bound by hydrogen bonding. Not surprisingly, no significant endoglucanase activity change is observed towards uniform avicel microcrystalline substrate (data not shown). Likewise, surfactants has been mixed right before addition of enzymes or at the beginning of the reaction to alter the microcrystalline substrate surface or enzyme confirmation.^33^ Preincubation with surfactants has been applied as pretreatment of untreated biomass sources or lignin containing substrates.^4^ Moreover, azoTAB-vis, azoTAB-UV, SDS, SDBS and DTAB added solutions at 37 °C towards avicel substrate are tested to understand if the activity enhancement of endoglucanase is temperature dependent. Almost identical relative activity increases are obtained at 37 °C compared to 25 °C (data not shown).

The optimal surfactant concentrations of 0.4 mM azoTAB under visible (1.5 mM) and UV light (7.5 mM);^14^ and 1.5 mM SDS (8 mM),^35^ SDBS (2.5 mM)^36^ and DTAB (15mM)^37^ are selected for the following experiments. The CMC (critical micelle concentration) points are specified in parenthesis. All of the conditions will characterize surfactant monomer and enzyme-substrate interactions rather than micelle and enzyme-substrate association.

In situ light illumination is shown in Figure 2a and 2b towards 4-nitrophenyl-*β*-D-cellobioside model substrate as opposed to avicel microcrystalline substrate in Figure 1. The addition of pre-converted azoTAB-visible and azoTAB-UV solutions lead to 2-fold (i.e. 1.0 nmol/s) and 3.8-fold (i.e. 1.9 nmol/s) reaction rate compared to the pure enzyme hydrolysis rate (i.e. 0.5 nmol/s). Endoglucanase activity enhancement with azoTAB towards p-nitrophenol-based substrate is significantly higher than avicel microcrystalline substrate. The degree of activity enhancement of endoglucanase is substrate specific as reported by others,^31, 38^ and our study also shows the difference between a soluble artificial substrate and an insoluble natural substrate. Nearly identical reaction velocities were obtained with in situ light illumination from visible to UV light and in reverse order. Additionally, SDS, SDBS and DTAB surfactant addition on endoglucanase activities towards the model substrate led to 2-3-fold relative activity increases (not shown). Again, this proves the specific relative activity increases by surfactants are substrate dependent.

**Figure 2.**
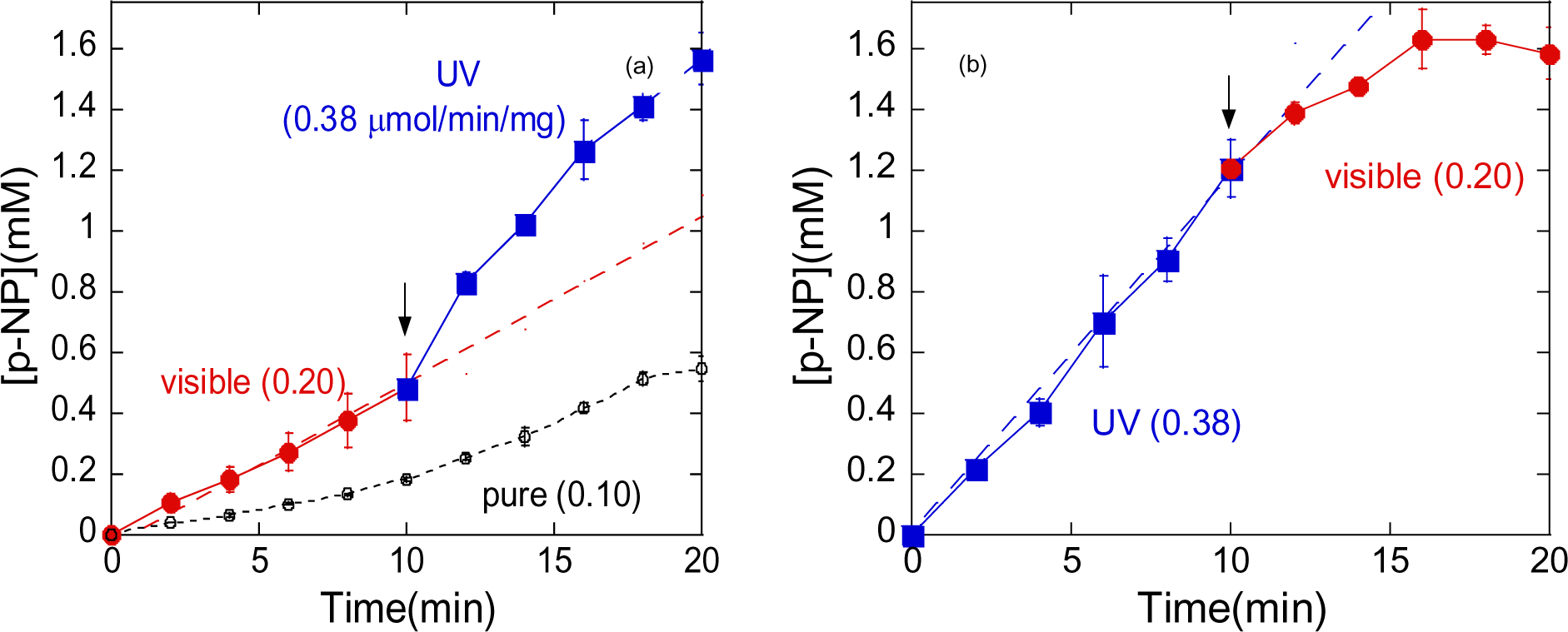
Light-responsive p-nitrophenol enzymatic cleavage product release from 4-nitrophenyl β-D-cellobioside substrate (5 mM) with 0.4 mM azoTAB and 0.3 mg/mL endoglucanase (a) visible to UV light exposure, (b) UV to visible light exposure. pH = 5, T = 37 °C. In situ light changes are pointed by arrows. The reaction rates are written in parenthesis. For reference, the product concentration in the presence of 0.4 mM azoTAB visible and UV light versus time are drawn with dashed lines.

Figure 3 shows the azoTAB addition to avicel saccharification compared to the pure enzyme condition with and without addition of 200 mM glucose to investigate enhanced endoglucanase activity visually. The majority of breaking of avicel with endoglucanase happens in the first three hours and then it slows down as displayed in the images. This is mainly due to the irreversible binding of endoglucanase onto the substrate surfaces and end-product inhibition as we discussed above. When we compare 3h pictures of pure endoglucanase and 0.4 mM azoTAB containing endoglucanase, less insoluble crystalline structures are seen in azoTAB containing reaction. When all time points are compared, less solid particles or improved cellulose saccharification is observed with the addition of azoTAB to endoglucanase enzyme solutions. Cellulose saccharification is a very slow process and it requires constant enzyme loading as we discussed above;^2^ thus, it is not surprising that we still see microcrystalline substates even at the end of 6^th^ day especially no additional enzyme are added in this experiment.

**Figure 3.**
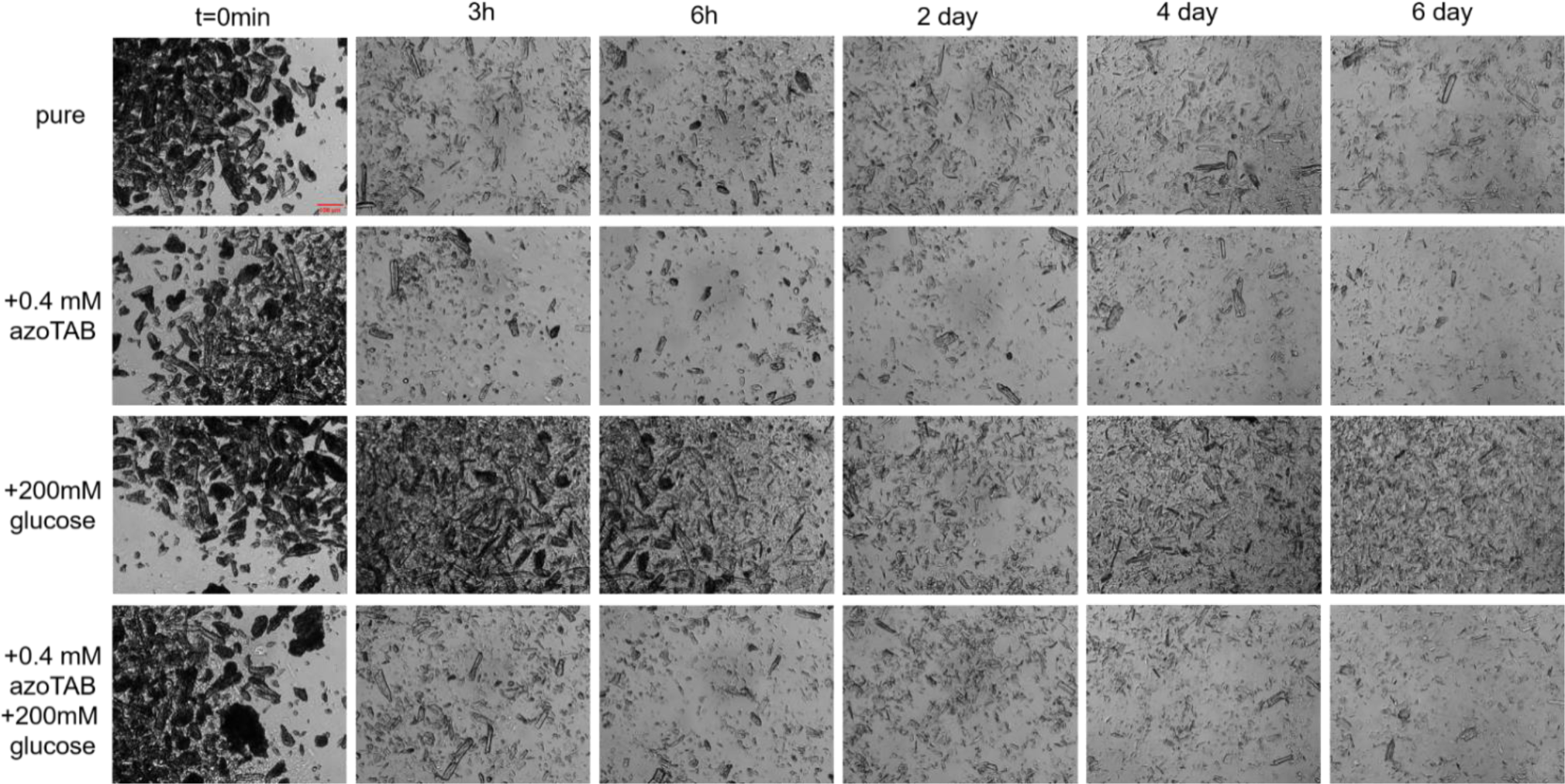
Optical microscopy images of avicel as a function of time. AzoTAB surfactant effect on avicel hydrolysis is observed with and without end-product glucose inhibitor. The scale bar shows 100 *µ*m.

End-product inhibition is another main concern of cellulose degradation and Figure 3 shows 200 mM added glucose effect on this process. It can be clearly seen that the solid content of 2^nd^-6^th^ day images of 200 mM glucose added reactions are more than 3h image of pure enzyme reaction. This simple comparison shows end-product inhibition of cellulose saccharification clearly. As a comparison, images of 0.4mM azoTAB containing 200 mM glucose added reactions show the improvement of the avicel degradation in the presence of end-product compared to the pure condition. Additionally, all the pictures shows that 200 mM glucose added pure enzyme solution has significantly more solid content than 0.4 mM azoTAB containing 200 mM glucose added endoglucanase solution. This image comparison can be concluded with the necessity of using surfactants in cellulose saccharification for modification of the solid substrate surface and decreasing of enzyme irreversible binding.

Insoluble crystalline cellulose has limited accessibility to soluble enzymes, and endoglucanase breaks random *β*-1,4-glycosidic bonds at the solid-liquid interface. Soluble endoglucanase adsorbs onto the surface of the cellulosic substrate and diffuse inside through pores and then breaks cellulose into insoluble and soluble sugars. As amphiphilic molecules, surfactants have a tendency to adsorb onto surfaces that provides modification on the cellulose surfaces as we discussed above, effect of azoTAB and other surfactants on endoglucanase adsorption on avicel is assessed. Adsorbed enzyme versus total enzyme concentration plot (Figure 4) shows that endoglucanase followed a Langmuir-type adsorption pattern with the assumptions of homogenous adsorption sites, reversible binding, one molecule binds to one adsorption site and the solutes on the surface do not interact with each other. As the endoglucanase concentration increases, the adsorbed protein on the solid surface raises. AzoTAB-vis and azoTAB-UV show 40-60 % higher amount of enzyme adsorption compared to the pure endoglucanase reaction. The adsorbed enzyme percentage increase with azoTAB can be associated to specific activity increases in Figure1 (30% for azoTAB-vis and 45% for azoTAB-UV). Similarly, the rate of cellulose hydrolysis is correlated to the adsorbed enzyme onto the substrate surface. ^19, 33^ The binding affinity is specific to enzyme properties and it was reported only insignificant amounts of BSA or β-glucosidase are adsorbed onto cellulose substrate.^17, 19^ Enzyme adsorption on solid substrate is affected by features of the enzyme and cellulose such as concentration, temperature, pH, structural properties or crystallinity of substrate, presence of inhibitors or enhancers.^4, 19^ Tyrosine (Tyr), tryptophan (Trp), glutamic acid (Glu) and aspartic acid (Asp) amino acids on the surface of cellulase also contributes to cellulase binding to the substrate.^7^ The cellulose binding affinity is modified with single amino acid substitutions; specifically tyrosine to tryptophan mutagenesis increases the binding affinity.^39^ The high adsorbed endoglucanase on avicel can be associated with special azoTAB interaction to tyrosine and tryptophan aromatic amino acids and azobenzene through π-π stacking in the binding sites. Overall, the adsorption results are correlated with relative hydrolysis rate for azoTAB effect results although 10-20% decrease in enzyme adsorption was observed for SDS, SDBS and DTAB containing samples in this study.

**Figure 4.**
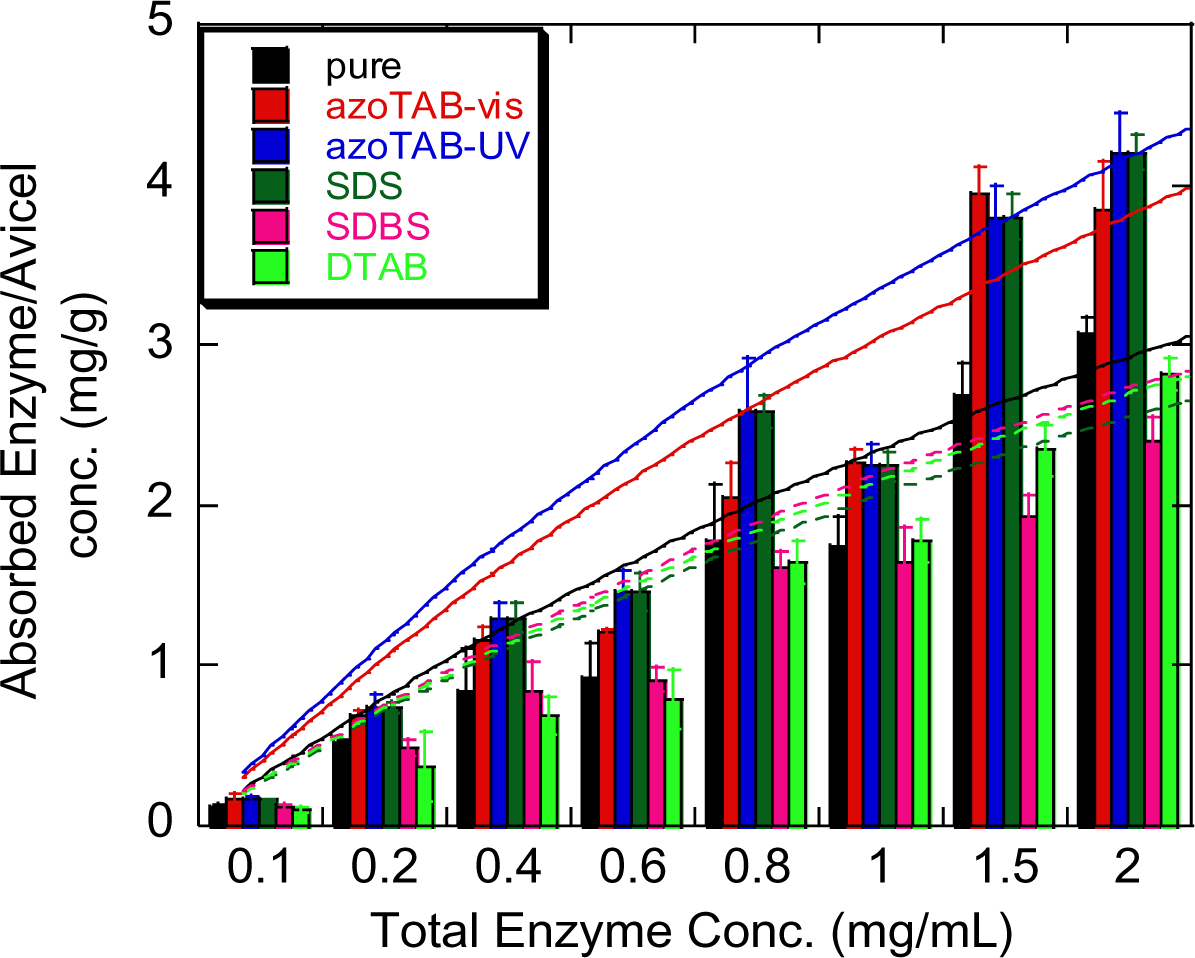
Adsorption isotherm of endoglucanase from *Aspergillus niger* on avicel (20mg/mL) in addition of azoTAB (0.4 mM) under visible and UV light, SDS (1.5 mM), SDBS (1.5mM) and DTAB (1.5 mM) surfactants (pH 5, T=25 °C). 30 min incubation time was used. The lines are drawn according to the individual fit of Langmuir isotherm to obtain *E*_max_ and *K*_ad_ values.

The linearized Langmuir isotherm equation (equation 2) is used to calculate maximum adsorbed enzyme concentration *E*_max_ and adsorption constant *K*_ad_ parameters from [*S*]/*E*_B_ versus 1/*E*_F_ plot in Figure S2 and the calculated parameters are listed in Table 1. The maximum enzyme concentration *E*_max_ increases from 7.89 mg enzyme/g substate to 11.52 mg enzyme/g substate (46% increase) and 12.92 mg enzyme/g substate (64% increase) with 0.4 mM azoTAB addition under visible and UV light, respectively. Only 3-10% increase of maximum enzyme adsorption is determined for SDS, SDBS and DTAB containing reactions while the adsorption constant or dissociation constant (*K*_ad_), as a reciprocal term for enzyme adsorption affinity decreases 22-30% for the surfactants that we used in this study. The endoglucanase activity improvement of SDS, SDBS and DTAB containing solutions is associated to dissociation constant or substrate modification while azoTAB addition to endoglucanase solution shows combination of the increase of endoglucanase adsorption on cellulose and substrate modification. Similar to our study, increase of *E*_max_ and decrease of *K*_ad_ are correlated with catalytic activity enhancement with different pretreatment conditions.^17^

**Table 1.** Comparison of adsorption parameters of endoglucanase on avicel (20 mg/mL) with and without azoTAB, SDS, SDBS and DTAB surfactants. *E*_max_ and *K*_ad_ values are calculated with Langmuir isotherm equation (equation 1 and 2) and the data points with the fits are represented in Figure 4 and Figure S2.

| | $E_{\max}(\text{mg/g}$<br>substrate) | $K_{\text{ad}}(\text{mg}$<br>enzyme/mL) | $R^2$ |
| --- | --- | --- | --- |
| pure | 7.89±0.89 | 0.33±0.06 | 0.9998 |
| azoTAB-vis | 11.52±0.18 | 0.27±0.10 | 0.9996 |
| azoTAB-UV | 12.92±1.13 | 0.26±0.04 | 0.9999 |
| SDS | 8.12±0.76 | 0.27±0.03 | 0.9998 |
| SDBS | 8.38±0.93 | 0.28±0.04 | 0.9998 |
| DTAB | 8.68±0.10 | 0.24±0.01 | 0.9999 |

Although Michaelis-Menten kinetics is an oversimplified method to analyze two-phase enzymatic reactions, it has been still a common applied model to identify catalytic constants and provide comparative results.^40^ The azoTAB effect on kinetic parameters, Michaelis constant (*K*_M_) and maximum velocity (*V*_max_), of endoglucanase is determined using Michaelis-Menten non-linear regression method (Figure 5) and listed in Table2. The turnover number, *k*_cat_ increases by 36% and 39% with azoTAB from 0.00033 s^-1^ to 0.00045s^-1^ and 0.00046s^-1^ under visible and UV light, respectively. Michaelis-Menten constant, *K*_M_ as a reciprocal term, lowers 3% and 30% with addition of 0.4 mM azoTAB under visible and UV light, respectively. Although, *k*_cat_ increase with addition of azoTAB under visible and UV light are similar, *K*_M_ parameters are quite different. The more hydrophilic UV photoisomer (45% activity enhancement) addition to endoglucase increases substrate affinity compared to the visible azoTAB (30% activity enhancement) due to the different increases the affinity. Overall, the total effect of azoTAB to catalytic efficiency (*k*_cat_/*K*_M_) of endoglucanase for avicel shows ∼40% and ∼100% enhancement from 0.03082 L*mg^-1^s^-1^ to 0.04322 L*mg^-1^s^-1^ and 0.06145 L*mg^-1^s^-1^ under visible and UV light, respectively. Hanes-Woolf plot is also used to acquire enzyme kinetics terms and similar results to nonlinear regression values are obtained.

**Figure 5.**
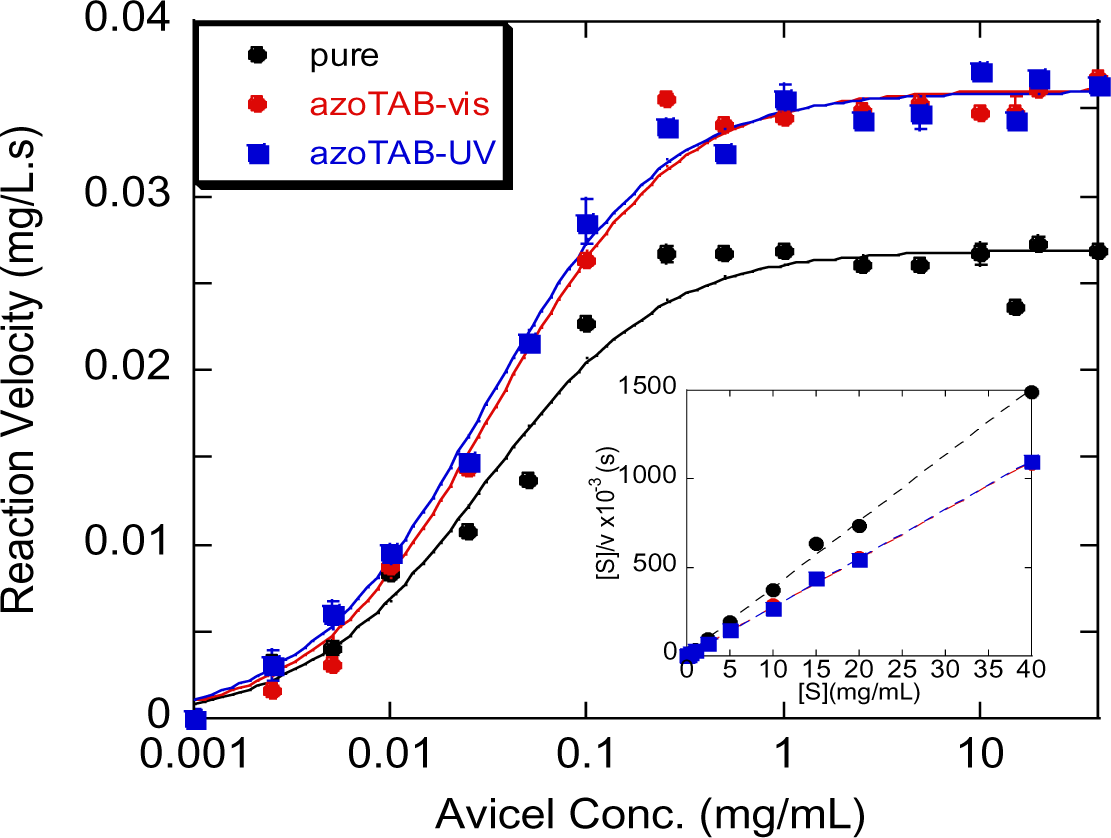
Reaction rates are acquired as a function of avicel microcrystalline substrate concentration in the presence of azoTAB (0.4 mM) under visible and UV light. Solid curves display the nonlinear regression fits of Michaelis-Menten equation. The inlet graph displays Hanes-Woolf plot, and the dashed lines show the fits to calculate kinetics constants.

**Table 2.** Reaction kinetics parameters of nonlinear regression fits of Michaelis-Menten equation and linear fit of Hanes-Woolf equation. The data points and fits are represented in the Figure 5.

|  | nonlinear regression |  |  | Hanes-Woolf constants |  |  |
| --- | --- | --- | --- | --- | --- | --- |
| | $k_{\text{cat}}(\text{s}^{-1}) \times 10^4$ | $K_{\text{M}}(\text{mg/L})$ | $k_{\text{cat}}/K_{\text{M}} (\text{L/mg.s})$ | $k_{\text{cat}}(\text{s}^{-1}) \times 10^4$ | $K_{\text{M}}(\text{mg/L})$ | $k_{\text{cat}}/K_{\text{M}} (\text{L/mg.s})$ |
| pure | 3.3 | 0.01071 | 0.03082 | 2.7 | 0.10425 | 0.00256 |
| azoTAB-vis | 4.5 | 0.01041 | 0.04322 | 3.7 | 0.12446 | 0.00294 |
| azoTAB-UV | 4.6 | 0.00748 | 0.06145 | 3.6 | 0.06884 | 0.00528 |

Fractal kinetic models have been applied to understand further information for the surfactant effect on enzyme-cellulose surfaces at the heterogeneous reaction. Enzymatic hydrolysis in time is fitted into equation (3)^20, 21^ to determine fractal parameters and kinetic constants for different surfactant solutions and the control (Table 3). Rate constant *k* is correlated with the initial rate of the enzyme concentration as well as the substrate accessibility while fractal exponent *h* gives the time effect of diminished rate.^21^ An addition of azoTAB leads to an increase of rate constants while conventional surfactant inclusion causes a slight decrease in the rate constants. This suggests slightly high surfactant concentrations (1.5 mM) altered endoglucanase confirmation in a way to decrease enzyme activity. At the same time, reciprocal term fractal exponent *h* of all surfactants added solutions lessened compared to the control samples. The combination of higher rate coefficient and lower fractal exponent shows the agreement on the optimum azoTAB concentrations of 0.4 mM both under visible and UV light. The results also indicate the optimal SDS, SDBS and DTAB concentrations were at 1.5 mM range, the endoglucanase activity is dropped at the further surfactant concentrations (Figure 1).

**Table 3.** Fractal kinetics parameter of endoglucanase on avicel (0.01 mg/mL) with and without surfactants. Fractal exponent and kinetics parameters are calculated with equation (3)^20, 21^ with 62.5 µg/mL endoglucanase enzyme towards 0.01mg/mL avicel.

| | $h$ | $k(\times 10^6)$ |
| --- | --- | --- |
| pure | 0.405 | 3.842 |
| azoTAB-vis | 0.329 | 4.380 |
| azoTAB-UV | 0.330 | 4.683 |
| SDS | 0.314 | 3.679 |
| SDBS | 0.301 | 3.270 |
| DTAB | 0.323 | 3.572 |

Accumulation of glucose and cellobiose inhibits endoglucanase catalytic activity^3^ due to the fact that the adsorption of cellulases declines and hence cellulase catalytic activity decreases as the solid content increases.^41^ Product inhibition is correlated with solid content increase rather than end product binding to catalytic sites of the enzyme or substrate.^42^ In here, Somogyi-Nelson method could not be applied due to the fact that high concentrations of end-products caused saturation of the measured dye. The reaction rate changes are minute for 50х diluted samples and hence making a judgement is trivial. Endoglucanase enzyme adsorption on the solid substrate as a function of inhibitor concentrations has been studied in the literature.^43, 44^ In order to understand the surfactant effect on endoglucanase adsorption, and thus; cellulose hydrolysis rate at high end-product concentrations, adsorption assay is studied at constant cellulose (20mg/mL) and endoglucanase (1 mg/mL) concentrations with 0-200 mM glucose and cellobiose. Cellobiose is found to be a stronger inhibitor compared to glucose at the same concentrations in Figure 6 similar to the earlier study.^42^ Again, this proves the solid content and cellulase activity since molecular weight of cellobiose is twice of glucose. In a similar fashion, adsorbed endoglucanase amount is found almost identical for 50 mM cellobiose-100 mM glucose and 100 mM cellobiose-200 mM glucose. The relative increment of enzyme activity by addition of all surfactants are similar throughout different inhibitor concentrations. For example, BSA is used to understand the solid content effect, and 1 mg/mL BSA addition decreases 60% endoglucanase activity as well as surfactant-added enzyme catalytic activity (data is not shown).

**Figure 6.**
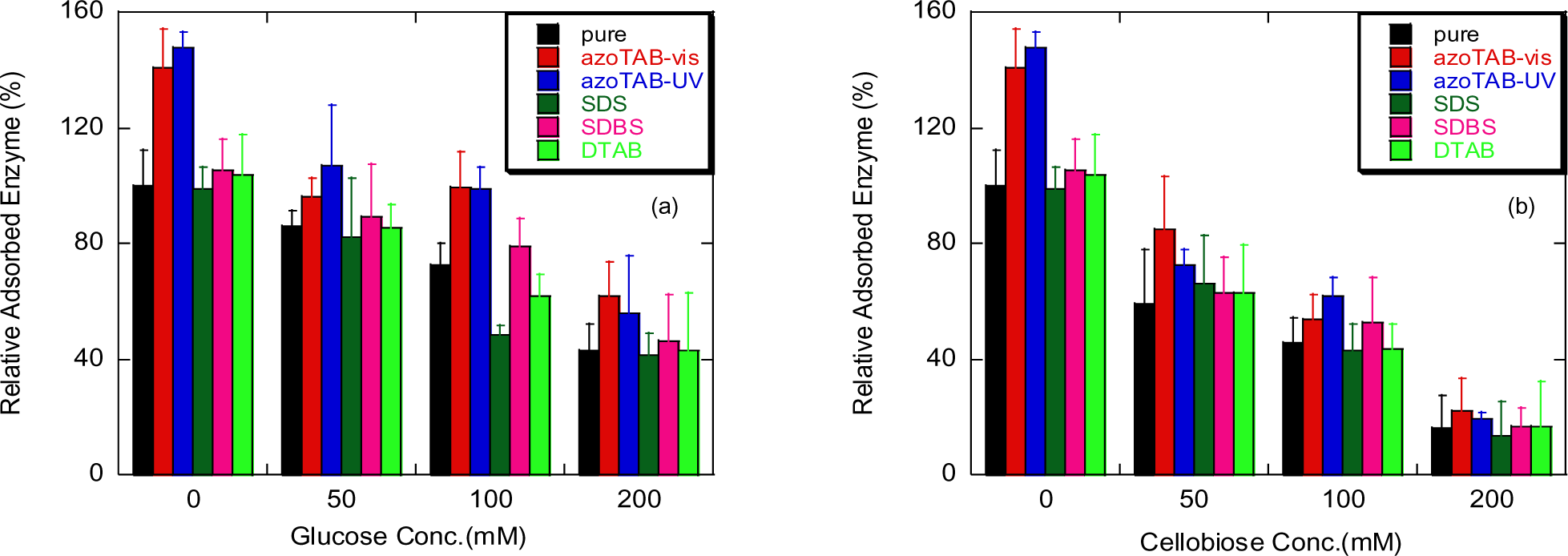
Percentage of the adsorbed endoglucanase (1 mg/mL) on avicel (20 mg/mL) as a function of (a) glucose and (b) cellobiose end-products with and without azoTAB (0.4 mM), SDS (1.5 mM), SDBS (1.5 mM) and DTAB (1.5 mM) surfactants.

The fusion protein of β-glucosidase and endoglucanase is designed and applied for carboxymethyl cellulose hydrolysis.^40^ Endoglucanase and cellobiohydrolase mixtures have always been used synergistically towards microcrystalline cellulose avicel.^23, 45^ Therefore, four different enzyme mixtures of endoglucanase; (1) only endoglucanase, (2) endoglucanase and β-glucosidase, (3) endoglucanase and cellobiohydrolase, and (4) all three enzymes of endoglucanase, β-glucosidase and cellobiohydrolase have selected to study the effect of surfactants on synergistic activity of cellulase enzyme mixtures for initial and long-term behavior in Figure 7. As the reaction time increases, the reaction rate slows down as shown above in Figure 3 and reported in the literature.^46^ AzoTAB-UV addition shows the highest enzyme activity enhancement (i.e. 25-55% of catalytic activity increase) at four different enzyme formulations and different time points. While 6-15 % light-responsive relative activity is observed throughout endoglucanase and a mixture of endoglucanase and β-glucosidase at different time points, the light-responsivity is the lowest at cellobiohydrolase enzyme containing solutions. Cellobiohydrolase addition lowers the enzyme activity enhancement with azoTAB surfactant both under visible and UV light compared to the pure enzyme solution. Similarly, poly ethylene glycol (PEG) addition increases cellobiohydrolase activity while shows no effect on endoglucanase activity,^23^ and Tween 20 increases cellulase adsorption while it shows no effect on β-glucosidase activity.^47^ It is also reported that while Tween 80 is an activator for cellobiohydrolase, it behaves as an inhibitor for endoglucanase or β-glucosidase.^48^ Presence of 0.4 mM azoTAB becomes more pronounced with the addition of β-glucosidase with 5-25% relative catalytic activity improvement of the enzyme mixtures. Likewise, our previous study reports 30% and 60% relative activity increase in β-glucosidase activity at 0.4 and 0.75 mM azoTAB addition under UV light, respectively.^14^ Our current results show the enhancement of β-glucosidase activity seemed conserved in the cellulase enzyme mixtures. Activity of β-glucosidase can be increased with addition of azoTAB either isolated β-glucosidases^14^ or mixtures of endoglucanases and cellohydrolases as shown in Figure 7. Overall, 50% of activity improvement is obtained with the addition of 0.4 mM of azoTAB independent of light wavelength for the combination of all three enzymes.

**Figure 7.**
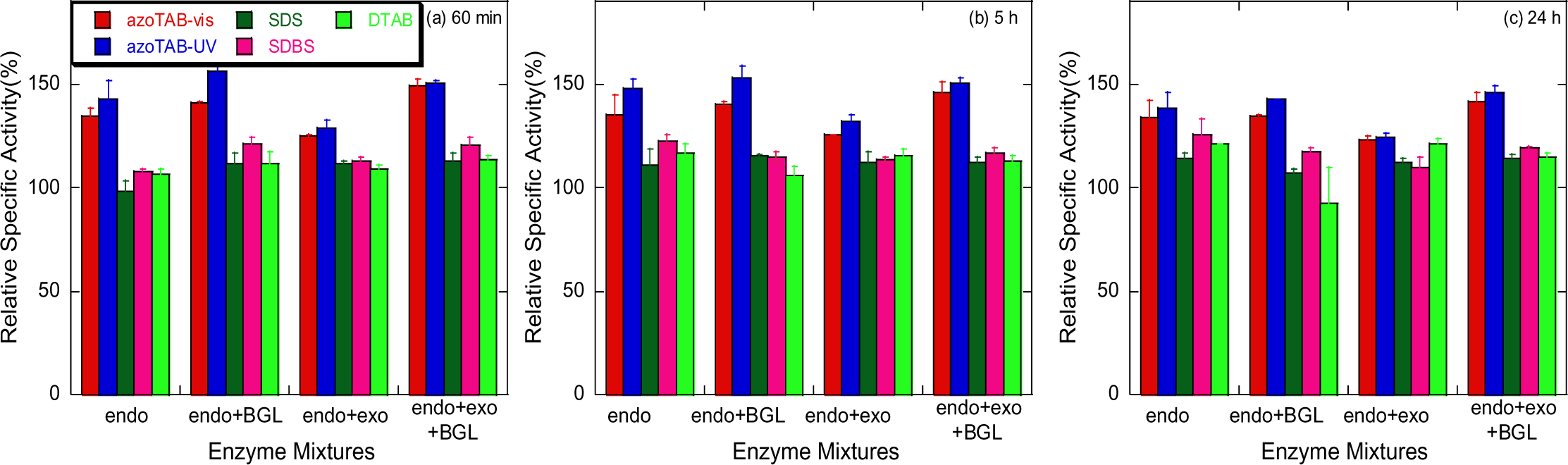
Relative activity of various enzyme mixtures of endoglucanase (62.5 μg/mL), cellobiohydrolase (120 μg/mL) and β-glucosidase (30 μg/mL) with addition of azoTAB (0.4 mM under visible or UV light), SDS (1.5 mM), SDBS (1.5 mM) and DTAB (1.5 mM) towards avicel (0.01 mg/mL), pH = 5, T = 25 °C. (a) Initial, (b) and (c) long-term relative saccharification rate of avicel.

On the other hand, cellobiohydrolase addition shows minimal change for other surfactants while β-glucosidase addition slightly increased the activity enhancement. Conventional surfactant addition causes activity increment in long term (i.e. 5h and 24h) while azoTAB conserved the constant activity enhancement of enzymes. Same activity enhancement is observed throughout 24 h for azoTAB while effect of other surfactants is seen in longer time. This could be explained with the main reason of surfactant addition into the cellulose degradation: (1) making the substrate surface more accessible to cellulase enzymes, (2) lowering irreversible binding of enzyme to substrate.^6, 30^ The explanation of why common surfactants shows activity rate improvement in long-term could be the differences of surfactant concentrations with azoTAB (azoTAB is 0.4 mM while other common surfactants are at 1.5 mM.) All in all, azoTAB has been shown to significantly improve (25-50% increment) both the initial rate and the long-term enzyme reaction rate in cellulose degradation.

## Conclusion

The catalytic activity and kinetics of endoglucanase at the solid-liquid interface have been investigated with of azoTAB photosurfactant under visible and UV light and compared to common surfactants. The enzyme specific activity is enhanced 45% with the addition of 0.4 mM azoTAB under UV light. Light-induced endoglucanase activity range becomes narrower (∼15%) for natural substrate avicel while it can be controlled between 2 to 4-fold toward p-nitrophenol-based model substrate (4-nitrophenyl β-D-cellobioside). The underlying reason of the 45% specific activity enhancement is mainly correlated with ∼40-50% increased adsorbed enzyme content and catalytic enzyme efficiency. In comparison, 5-10% slight catalytic activity increase was observed in addition of sodium dodecyl sulfate (SDS), sodium dodecyl benzene sulfonate (SDBS) and dodecyl trimethyl ammonium (DTAB) traditional surfactants and correlated with substrate affinity modification. Lastly, azoTAB addition maintained 45-50% enzyme catalytic rate increase on all three cellulase enzyme mixture of endoglucanase, cellobiohydrolase and β-glucosidase both for initial rate and long-term behavior. Overall, azoTAB could be applied as a promising activator of cellulase enzymes for biomass hydrolysis.

## Table of Content

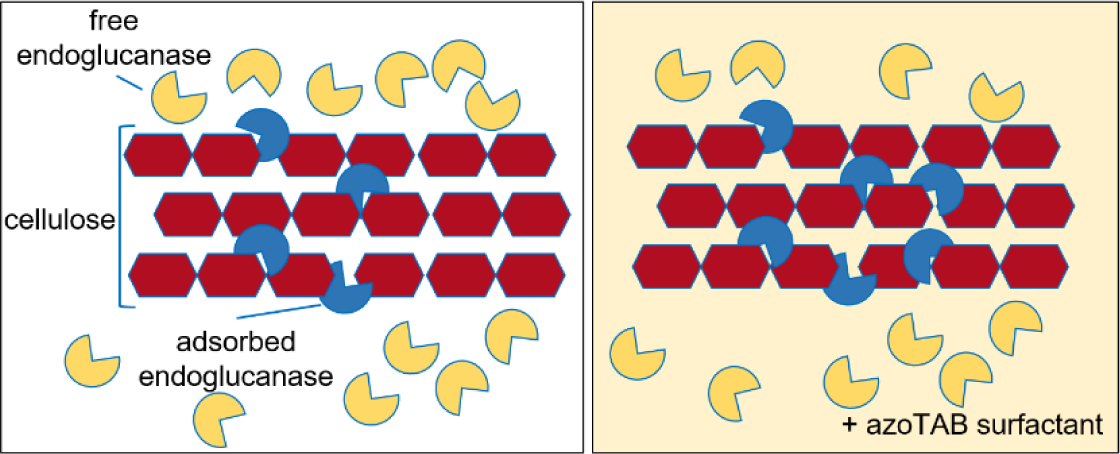

## Acknowledgement

This material is based upon work supported by the National Science Foundation under Grant 1758225. Any opinions, findings, and conclusions or recommendations expressed in this material are those of the author(s) and do not necessarily reflect the views of the National Science Foundation. The authors acknowledge Dr. Fariborz Nasertorabi of USC’s Bridge Institute for his assistance with enzyme purification studies.

## Notes

### Competing Interest Statement

The authors have declared no competing interest.

## References

(1) Mansouri, A.; Rihani, R.; Laoufi, A. N.; Özkan, M. Production of bioethanol from a mixture of agricultural feedstocks: Biofuels characterization Fuel 2016, 185, 612–621.

(2) Zhang, X.-Z.; Zhang, Y.-H. P. Cellulases: Characteristics, Sources, Production and Applications. In Bioprocessing Technologies in Biorefinery for Sustainable Production of Fuels, Chemicals, and Polymers, Yang, S.-T., El-Enshasy, H. A., Thongchul, N. Eds.; John Wiley & Sons, 2013; pp 131–146.

(3) Teugjas, H.; Valjamae, P. Product inhibition of Cellulases Studied with 14C-labeled Cellulose Subsrates. Biotechnology for Biofuels 2013, 6: 104.

(4) Qing, Q.; Yang, B.; Wyman, C. E. Impact of surfactants on pretreatment of corn stover. Bioresor. Technol. 2010, 101(15), 5941–5951. DOI: 10.1016/j.biortech.2010.03.003.

(5) Rastegari, A. A.; Bordbarb, A.-K.; Taheri-Kafranib, A. Interaction of cellulase with cationic surfactants: Using surfactant membrane selective electrodes and fluorescence spectroscopy Colloids and Surfaces B: Biointerfaces 2009, 73, 132–139

(6) Eriksson, T.; Borjesson, J.; Tjerneld, F. Mechanism of surfactant effect in enzymatic hydrolysis of lignocellulose Enzyme and Microbial Technology 2002, 31 (353–364).

(7) Eckard, A. D.; Muthukumarappan, K.; Gibbons, W. A Review of the Role of Amphiphiles in Biomass to Ethanol Conversion Applied Sciences 2013, 3, 396–419.

(8) Seo, D.-J.; Fujita, H.; Sakoda, A. Structural changes of lignocelluloses by a nonionic surfactant, Tween 20, and their effects on cellulase adsorption and saccharification. Bioresource Technology 2011, 102, 9605–9612.

(9) Zhang, J.; Lee, C. T. J. Controlling Membrane Protein Folding with Light Illumination and Catanionic Surfactant Systems. University of Southern California, 2008.

(10) Mirarefi, P.; Lee Jr., C. T. Photo-induced Unfolding and Inactivation of Bovine Carbonic Anhydrase in the Presence of a Photoresponsive Surfactant. Biochimica et Biophysica Acta 2010, 1804, 106–114.

(11) Hamill, A. C.; Wang, S. C.; Lee, C. T. Probing lysozyme conformation with light reveals a new folding intermediate. Biochemistry 2005, 44 (46), 15139–15149. DOI: 10.1021/bi051646c.

(12) Lee, J. C. T.; Smith, K. A.; Hatton, T. A. Photocontrol of Protein Folding: The Interaction of Photosensitive Surfactants with Bovine Serum Albumin. Biochemistry 2005, 44, 524–536.

(13) Seidel, Z. P. Enhancement of Biofuel Enzyme Activity and Kinetics with AzoTAB Surfactants. University of Southern California, University of Southern California, 2021. https://digitallibrary.usc.edu/Share/47kkx3q62s5i1uu04t53484r733k040o.

(14) Seidel, Z. P.; Lee, C. T. J. Enhanced Activity of the Cellulase Enzyme β-Glucosidase upon Addition of an Azobenzene-Based Surfactant. ACS Sustainable Chem. Eng. 2020, 8, 1751–1761.

(15) Hayashita, T.; Kurosawa, T.; Miyata, T.; Tanaka, K.; Igawa, M. Effect of structural variation within cationic azo-surfactant upon photoresponsive function in aqueous solution. Colloid & Polymer Science 1994, 272, 1611–1619.

(16) Nelson, N. A photometric adaptation of the Somogyi method for the determination of glucose. J. Biol. Chem. 1944, 153, 375–380.

(17) Bommarius, A. S.; Katona, A.; Cheben, S. E.; Patel, A. S.; Ragauskas, A. J.; Knudson, K.; Pu, Y. Cellulase Kinetics as a Function of Cellulose Pretreatment. Metabolic Engineering 2008, 10, 370–381.

(18) Kadam, K. L.; Rydholm, E. C.; McMillan, J. D. Development and Validation of a Kinetic Model for Enzymatic Saccharification of Lignocellulosic Biomass. Biotechnol. Prog. 2004, 20, 698–705.

(19) Lee, S. B.; Shin, H. S.; Ryu, D. D. Y. Adsorption of Cellulase on Cellulose: Effect of Physicochemical Properties of Cellulose on Adsorption and Rate of Hydrolysis. Biotechnology and Bioengineering 1982, XXIV, 2137–2153. DOI: 10.1002/bit.260241003

(20) Wang, Z.; Feng, H. Fractal kinetic analysis of the enzymatic saccharification of cellulose under different conditions. Bioresource Technology 2010, 101, 7995–8000. DOI: 10.1016/j.biortech.2010.05.056.

(21) Wang, Z.; Xu, J.-H.; Feng, H.; Qi, H. Fractal Kinetic Analysis of Polymers/Nonionic Surfactants to Eliminate Lignin Inhibition in Enzymatic Saccharification of Cellulose. Bioresource Technology 2011, 102, 2890–2896.

(22) Maurer, S. A.; Bedbrook, C. N.; Radke, C. J. Competitive Sorption Kinetics of Inhibited Endo-and Exoglucanases on a Model Cellulose Substrate. Langmuir 2014, 28, 14598–14608. DOI: 10.1021/la3024524.

(23) Hsieh, C.-w. C.; Cannella, D.; Jorgensen, H.; Felby, C.; Thygesen, L. G. Cellobiohydrolase and Endoglucanase Respond Differently to Surfactants During the Hydrolysis of Cellulose. Biotechnology for Biofuels 2015, 8:52.

(24) Vries, R. P.; Visser, J. Aspergillus Enzymes Involved in Degradation of Plant Cell Wall Polysaccharides. Microbiology and Molecular Biology Reviews 2001, 65 (4), 497–522.

(25) Lee, N. E.; Lima, M.; Woodward, J. Hydrolysis of Cellulose by a Mixture of Trichoderma reesei Cellobiohydrolase and Aspergillus niger Endoglucanase. Biochimica et Biophysica Acta 1988, 967, 437–440.

(26) Coral, G.; Arikan, B.; Unaldi, M. N.; Guvenmez, H. Some Properties of Crude Carboxymethyl Cellulase of Aspergillus niger Z10 Wild-Type Strain. Turk J. Biol. 2002, 26, 209–213.

(27) Wu, J.; Ju, L.-K. Enhancing Enzymatic Saccharification of Waste Newsprint by Surfactant Addition. Biotechnol. Prog. 1998, 14, 649–652.

(28) Stoner, M. R.; Dale, D. A.; Gualfetti, P. J.; Becker, T.; Randolph, T. W. Surfactant-Induced Unfolding of Cellulase: Kinetic Studies. Biotechnol. Prog. 2006, 22, 225–232. DOI: 10.1021/bp0501468.

(29) Madsen, J. K.; Pihl, R.; Moller, A. H.; Madsen, A. T.; Otzen, D. E.; Anderson, K. K. The anionic biosurfactant rhamnolipid does not denature industrial enzymes Front. Microbiol. 2015, 6: 292, 1–13. DOI: 10.3389/fmicb.2015.00292.

(30) Ooshima, H.; Sakata, M.; Harano, Y. Enhancement of Enzymatic Hydrolysis of Cellulose by Surfactant. Biotechnology and Bioengineering 1986, 28, 1727–1734.

(31) Sahin, S.; Ozmen, I.; Biyik, H. Industrial Applications of Endoglucanase Obtained from Novel and Native Trichoderma atroviride. Chem. Biochem. Eng. Q. 2016, 30 (2), 265–278.

(32) Ying, W.; Xu, Y.; Zhang, J. Surfactants protect the activities of endoglucanase and cellobiohydrolase from gas-liquid interface. Industrial Crops and Products 2021, 171. DOI: 10.1016/j.indcrop.2021.113958.

(33) Helle, S. S.; Duff, S. J. B.; Cooper, D. G. Effect of Surfactants on Cellulose Hydrolysis. Biotechnology and Bioengineering 1993, 42, 611–617.

(34) Kim, H. J.; Kim, S. B.; Kim, C. J. The effects of nonionic surfactants on the pretreatment and enzymatic hydrolysis of recycled newspaper. Biotechnology and Bioprocess Engineering 2007, 12, 147–151.

(35) Naskar, B.; Dey, A.; Moulik, S. P. Counter-ion Effect on Micellization of Ionic Surfactants: A Comprehensive Understanding with Two Representatives, Sodium Dodecyl Sulfate (SDS) and Dodecyltrimethylammonium Bromide (DTAB). J Surfact Deterg 2013, 16, 785–794.

(36) Hait, S. K.; Majhi, P. R.; Blume, A.; Moulik, S. P. A Critical Assessment of Micellization of Sodium Dodecyl Benzene Sulfonate (SDBS) and Its Interaction with Poly (vinyl pyrrolidone) and Hydrophobically Modified Polymers, JR 400 and LM 200. J. Phys. Chem B 2003, 107, 3650–3658.

(37) Bales, B. L.; Zana, R. Characterization of Micelles of Quaternary Ammonium Surfactants as Reaction Media I: Dodeclytrimethylammonium Bromide and Chloride. J. Phys. Chem. B 2002, 106, 1926–1939.

(38) Akram, F.; Haq, I. u.; Ikram, W.; Mukhtar, H. Insight Perspectives of Thermostable Endoglucanases for Bioethanol Production: A Review. Renewable Energy 2018, 122, 225–238.

(39) Linder, M.; Lindeberg, G.; Reinikainen, T.; Teeri, T. T.; Pettersson, G. The difference in affinity between two fungal cellulose-binding domains is dominated by a single amino acid substitution. FEBS Letters 1995, 372, 96–98. DOI: 10.1016/0014-5793(95)00961-8.

(40) Adlakha, N.; Sawant, S.; Anil, A.; Lali, A.; Yazdani, S. S. Specific Fusion of b-1,4-endoglucanase and b-1,4-glucosidase enhances celluloytic activity and helps in channeling of intermediates. Applied and Environmental Microbiology 2012, 78(20), 7447–7454. DOI: 10.1128/AEM.01386-12.

(41) Ogeda, T. L.; Silva, I. B.; Fidale, L. C.; Seoud, O. A. E.; Petri, D. F. S. Effect of cellulose physical characteristics, especially the water sorption value, on the efficiency of its hydrolysis catalyzed by free or immobilized cellulase. Journal of Biotechnology 2012, 157(1), 246–252. DOI: 10.1016/j.jbiotec.2011.11.018.

(42) Kristensen, J. B.; Felby, C.; Jorgensen, H. Yield-determining factors in high-solids enzymatic hydrolysis of lignocellulose. Biotechnology for Biofuels 2009, 2: 11, 1–10. DOI: 10.1186/1754-6834-2-11.

(43) Kristensen, J. B.; Borjesson, J.; Bruun, M. H.; Tjerneld, F.; Jorgensen, H. Use of surface active additives in enzymatic hydrolysis of wheat straw lignocellulose Enzyme and Microbial Technology 2007, 40, 888–895

(44) Hsieh, C.-w. C.; Cannella, D.; Jorgensen, H.; Felby, C.; Thygesen, L. G. Cellulase Inhibition by High Concentrations of Monosaccharides. J. Agric. Food Chem. 2014, 62, 3800–3805.

(45) Medve, J.; Karlsson, J.; Lee, D.; Tjerneld, F. Hydrolysis of microcrystalline cellulose by cellobiohydrolase I and endoglucanase II from Trichoderma reesei: Adsorption, sugar production pattern, and synergism of the enzymes. Biotechnology and Bioengineering 1998, 59(5), 621–634. DOI: 10.1002/(SICI)1097-0290(19980905)59:5<621::AID-BIT13>3.0.CO;2-C.

(46) Bansal, P.; Hall, M.; Realff, M. J.; Lee, J. H.; Bommarius, A. S. Modelling cellulase kinetics on lignocellulosic substrates. Biotechnology Advances 2009, 27, 833–848. DOI: 10.1016/j.biotechadv.2009.06.005.

(47) Seo, D.-J.; Fujita, H.; Sakoda, A. Effects of a non-ionic surfactant, Tween 20, on adsorption/desorption of saccharification enzymes onto/from lignocelluloses and saccharification rate. Adsorption 2011, 17, 813–822. DOI: 10.1007/s10450-011-9340-8.

(48) Xin, D.; Yang, M.; Chen, X.; Zhang, Y.; Ma, L.; Zhang, J. Improving the Hydrolytic Action of Cellulases by Tween 80: Offsetting the Lost Activity of Cellubiohydrolase Cel7A. ACS Sustainable Chem. Eng. 2017, 5, 11339–11345.

