## Supporting Information for "Improved Activity and Kinetics of Endoglucanase Biofuel Enzyme with Addition of an AzoTAB Surfactant"

Number of pages: 2

Number of figures: 2

Number of tables: 0

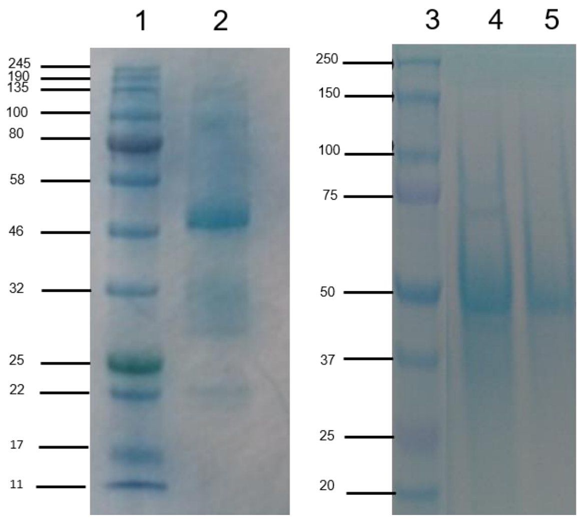


**Figure S1.** SDS-PAGE of crude cellulase (lane 2) and purified endoglucanase after anion exchange column Sephadex (lane 4 and 5) versus molecular weight protein ladders (lane 1 and 3)

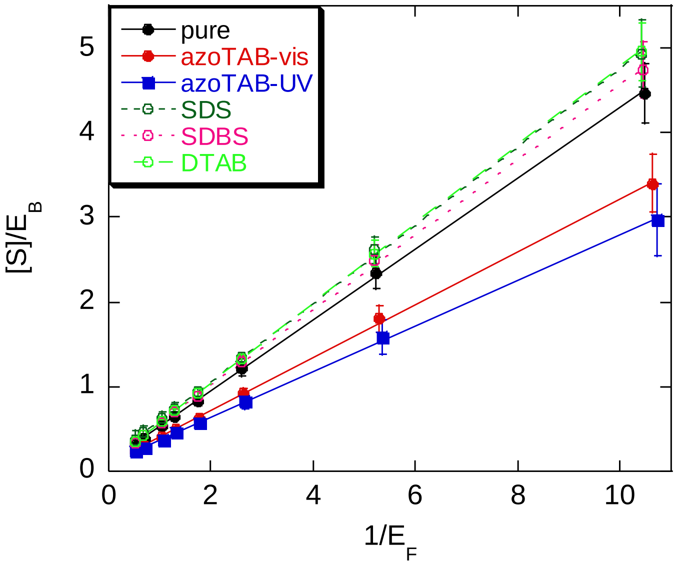


**Figure S2.** Linearized form of Langmuir adsorption model to calculate $E_{max}$ and $K_{ad}$ values (Table 2)
